# Social network analysis reveals context-dependent kin relationships in wild sulphur-crested cockatoos, *Cacatua galerita*

**DOI:** 10.1101/2022.04.06.486687

**Authors:** J. Penndorf, K. M. Ewart, B. C. Klump, J. M. Martin, L. M. Aplin

**Affiliations:** Cognitive and Cultural Ecology Research Group, Max Planck Institute of Animal Behavior, Radolfzell, Germany; Australian Museum Research Institute, Sydney, NSW, Australia; School of Life and Environmental Sciences, The University of Sydney, Camperdown, NSW, Australia; Institute of Science and Learning, Taronga Conservation Society Australia, Mosman, NSW, Australia; Centre for the Advanced Study of Collective Behaviour, University of Konstanz, Konstanz, Germany; Division of Ecology and Evolution, Research School of Biology, The Australian National University, Canberra, Australia

**Keywords:** *Cacatua galerita*, kinship networks, parrots, population genetics, social network analysis, urban ecology

## Abstract

1. A preference to associate with kin facilitates inclusive fitness benefits, and increased tolerance or cooperation between kin may be an added benefit of group living. Many species exhibit preferred associations with kin, however it is often hard to disentangle active preferences from passive overlap, for example caused by limited dispersal or inheritance of social position.
2. Many parrots exhibit social systems consisting of pair-bonded individuals foraging in variably sized fission-fusion flocks within larger communal roosts of hundreds of individuals. Previous work has shown that, despite these fission-fusion dynamics, individuals can exhibit long-term preferred foraging associations outside their pair-bonds. Yet the underlying drivers of these social preferences remains largely unknown.
3. In this study, we use a network approach to examine the influence of kinship on social associations and interactions in wild, communally roosting sulphur-crested cockatoos, *Cacatua galerita*. We recorded roost co-membership, social associations and interactions in 561 individually marked birds across three neighbouring roosts. We then collected genetic samples from 205 cockatoos, and conducted a relationship analysis to construct a kinship network. Finally, we tested correlations between kinship and four social networks: association, affiliative, low-intensity aggression, and high-intensity aggression.
4. Our result showed that while roosting groups were clearly defined, they showed little genetic differentiation or kin structuring. Between roost movement was high, with juveniles, especially females, repeatedly moving between roosts. Both within roosting communities, and when visiting different roosts, individuals preferentially associated with kin. Supporting this, individuals were also more likely to allopreen kin. However, contrary to expectation, individuals preferred to direct aggression towards kin, with this effect only observed when individuals shared roost membership.
5. By measuring social networks within and between large roosting groups, we could remove potential effects of passive spatial overlap on kin structuring. Our study reveals that sulphur-crested cockatoos actively prefer to associate with kin, both within and between roosting groups. By examining this across different interaction types, we further demonstrate that sulphur-crested cockatoos exhibit behavioural and context-dependent interaction rules towards kin. Our results help reveal the drivers of social association in this species, while adding to the evidence for social complexity in parrots.

## Introduction

Living in groups can provide a range of benefits to all individual members (Brown and Brown, 1987). These benefits can be further enhanced by preferentially associating with specific others (Krause et al., 2002). For example, associations with individuals sharing characteristics such as age (Aplin et al., 2021), sex (reviewed by Ruckstuhl, 2007), body size (Pitcher et al., 1986; Croft et al., 2005), or rank (Smith et al., 2007) can reduce conflict (Dadda et al., 2005) and predation risk (Sorato et al., 2012). Moreover, associations with kin have been shown to provide fitness benefits that range from higher reproductive success (Silk et al., 2009) and inclusive fitness (Hamilton, 1964; Grafen, 1984; Levin and Grafen, 2019, though see Birch, 2017) to reduced stress levels (Gerlach et al., 2007) and increased longevity (Silk et al., 2010). Conversely, associating with kin can be disadvantageous if such associations increase the risk of inbreeding, or lead to competition between relatives (Evans et al., 2021). However, these costs can be partly mitigated via context-dependent social decision-making, for example avoiding kin during the breeding season while preferring them at other times (Evans et al., 2021). Yet while the costs and benefits of associating with kin have been extensively explored, the underlying processes leading to such patterns warrants further research.

Association with kin might reflect active choice, but alternatively might arise from passive mechanisms such as limited dispersal (Klauke et al., 2016) or an inheritance of social position (Ilany et al., 2021). There are two likely avenues to help disentangle active and passive drivers for social associations. The first is to examine this question in social systems exhibiting fission-fusion social dynamics, where larger groups or communities frequently split into subgroups of variable composition. In systems with such fluid group membership, sex, age and breeding status have all been shown to affect associations, with individuals sharing these characteristics being more associated than expected by chance (Chiyo et al., 2011; Zeus et al., 2018; Aplin et al., 2021). When considering kinship however, the pattern is both more mixed and context-dependent. In species where individuals forms alliances, individuals jointly direction aggressions towards others tend to be more related than expected by chance (e.g. dolphins: Krützen et al., 2003; Parsons et al., 2003; monk parakeets: Dawson Pell et al., 2021). While a more general tendency to associate with kin has also been reported (e.g., Elephants, *Loxodonta africana*: Chiyo et al., 2011; Geese, *Branta leucopsis*: Kurvers et al., 2013; Giraffes, *Giraffa camelopardalis* : Carter et al., 2013), other species show no such preference; for example, roosting patterns in big brown bats (*Eptesicus fuscus*, Metheny et al., 2008) are not affected by kinship. Perhaps most intriguingly, a recent comparative study investigating kinship, social networks, and roost-switching behaviour in nine species of bats (Wilkinson et al., 2019), found that the preference to associate with kin was stronger in species with more roost-switching. This suggests a possible non-linear relationship between social structure and kinship, where the benefits of associating with kin may not only be elevated in highly structured societies (e.g., cooperative breeding systems), but also be higher in social contexts where very few individuals are known to one another.

A second potential avenue is to examine variation in interaction decisions while controlling for their social opportunity. Here the assumption is that if individuals can recognise kin and are preferentially associating with them, they should also be more likely to direct affiliative behaviours towards relatives (e.g. grooming and food sharing), and be less likely to engage in aggressive interactions (reviewed by Hepper, 1986; Waldman, 1988). Supporting this, a meta-analysis of grooming in primates found a general positive correlation with kinship, although it also supported a larger role for direct reciprocity (Schino and Aureli, 2010). While the literature on social interactions is extensive, most studies have examined these questions in groups with high relatedness (e.g. in cooperative breeders, Madden et al., 2012). There are relatively few examples of studies examining the influence of kinship on social decision-making in more open, fission-fusion species.

Here, we examined the influence of kinship on social associations and interactions in sulphurcrested (SC-) cockatoos, *Cacatua galerita*. Similar to many parrot species (Hardy, 1965; Power, 1966; Styche, 2000), SC-cockatoos live in large-scale fission-fusion societies, where stable night-roost aggregations of up to several hundred individuals (Lindenmayer et al., 1996) break-up during the day into smaller flocks that forage in a collective home range around the roost site. Foraging flocks have a fluid membership and are of variable size depending on activity, environment, and season (Noske et al., 1982; Aplin et al., 2021). There is little evidence of any kin structuring within roosts in SC-cockatoos or any other parrot species (Wright and Wilkinson, 2001, but see Blanco et al., 2021). In addition to within roost dynamics, between-roost movements are common in SC-cockatoos (Aplin et al., 2021), although their predictors are largely unknown. The combination between within- and between-roost social dynamics confer a large choice of potential interaction partners, and previous work in this species has shown that SC-cockatoos form non-random social associations, with stable, long-term bonds beyond the pair (Aplin et al., 2021). However, little is known about the factors underlying association preferences, or the predictors of social interactions such as allopreening or aggression.

Sulphur-crested cockatoos are urban persisters, and in such environments, populations are relatively habituated and often forage on the ground (Davis et al., 2017; Kirksey et al., 2018), giving a rare opportunity to conduct detailed behavioural observations. First, we habituated and marked 561 SC-cockatoos across five neighbouring roosting communities. We then collected genetic material for 393 individuals and performed an analysis of genetic relatedness for these five sites. Second, we collected detailed observations at three of these roost sites, and an intermediate regularly visited foraging site, as well as censusing roost co-membership. This resulted in 205 individuals where we had information on both kinship and social interactions. We then examined the correlation between kinship and sociality using social network analysis to compare kinship networks with social associations (foraging flock co-occurence) and three interaction networks - allopreening (an affiliative behaviour), and low- and high-intensity aggressive interactions. Finally, we performed a second analysis to examine behaviour at two scales: within and between roosting communities.

Previous work on this population of SC-cockatoos has revealed that individuals maintain long-term bonds outside of the pair, which presumably confer benefits to individuals (Aplin et al., 2021). Given the additional predicted benefits of interacting with kin, we predicted that these stronger bonds would often be with close kin. We further predicted that, if this represented an active decision (rather than, for example, delayed dispersal of offspring), SC-cockatoos would show a preference to associate with kin when visiting other roost groups, or when meeting kin from other roost groups at foraging sites. Finally, we predicted that individuals would preferentially direct affiliative interactions towards kin, and be less likely to direct aggressive interactions towards kin; we further predicted that this would hold when controlling for social opportunity. Our multi-step analytical approach removes the potential effect of spatial overlap and philopatry on any correlation between kinship and social associations, while allowing us to ask whether SC-cockatoos show behavioural or context-dependent social interaction rules towards kin that would infer both individual recognition and a degree of social complexity (Wilkinson et al., 2019).

## Methods

### Study system

Sulphur-crested cockatoos are large (700-1200g), slow breeding, long-lived parrots. They are common across a large native range in Australia and Papua New Guinea, and populations appear to be increasing, including in human-modified areas (Davis et al., 2017). In the Sydney population, the majority of individuals are non-breeding, forming stable communal roosts of tens to hundreds of individuals, usually in parkland with remnant old growth trees, similar to previous descriptions at other sites (Lindenmayer et al., 1996). Tree hollows for nesting are a limited resource, and breeding pairs (approximately 20-40% of adults), defend nearby nest hollows year-round, but often roost with the rest of the group when not actively breeding (*personal observation*). Most individuals are site faithful, with a previous study on this population finding that birds used an average of two roost sites over a period of five years (Aplin et al., 2021). Within roosting communities, SC-cockatoos exhibit fission-fusion dynamics during foraging, and individuals of different roosts associate when feeding on a variety of resources in the surrounding area (Aplin et al., 2021). In urban areas, resources include human derived and provisioned food (Kirksey et al., 2018; Klump et al., 2021), making birds relatively easy to habituate.

### Habituation, marking and morphometric data collection

Birds were habituated to human observers at suburban parks close to three focal roosts (Balmoral Beach BA, Clifton Gardens CG, and Northbridge Golf Club NB), and at the nearest two neighbouring roosts (Sydney Botanic Gardens BG and Ivanhoe Park MA; Figure 1a.) Over 2-3 weeks, individuals were attracted to the ground at the same time each day with small amounts of sunflower seed, a commercially available food commonly eaten by SC-cockatoos in urban and agricultural areas (Noske et al., 1982). The observer would remain as close as possible to the foraging flock until the birds began to forage on the ground around the observer and birds could be lightly touched on the back without eliciting an adverse response. Once birds were habituated, they were temporarily and non-invasively marked using non-toxic fabric dye (Marabu Fashion Spray, MARABU GMBH), with each individual receiving a unique mark of one to three colours placed on the middle of the back (Figure 1b). This method resulted in 536 uniquely marked individuals across the five roosts. In addition, 17 birds present in our study area had already been caught, ringed and wing-tagged as part of a previous study (Figure 1c, Davis et al., 2017, Aplin et al., 2021), and 8 birds were uniquely recognisable due to morphological variation (e.g., healed injuries). This resulted in 561 recognisable individuals, which we estimate to be >90% of the local population.

**Figure 1:**
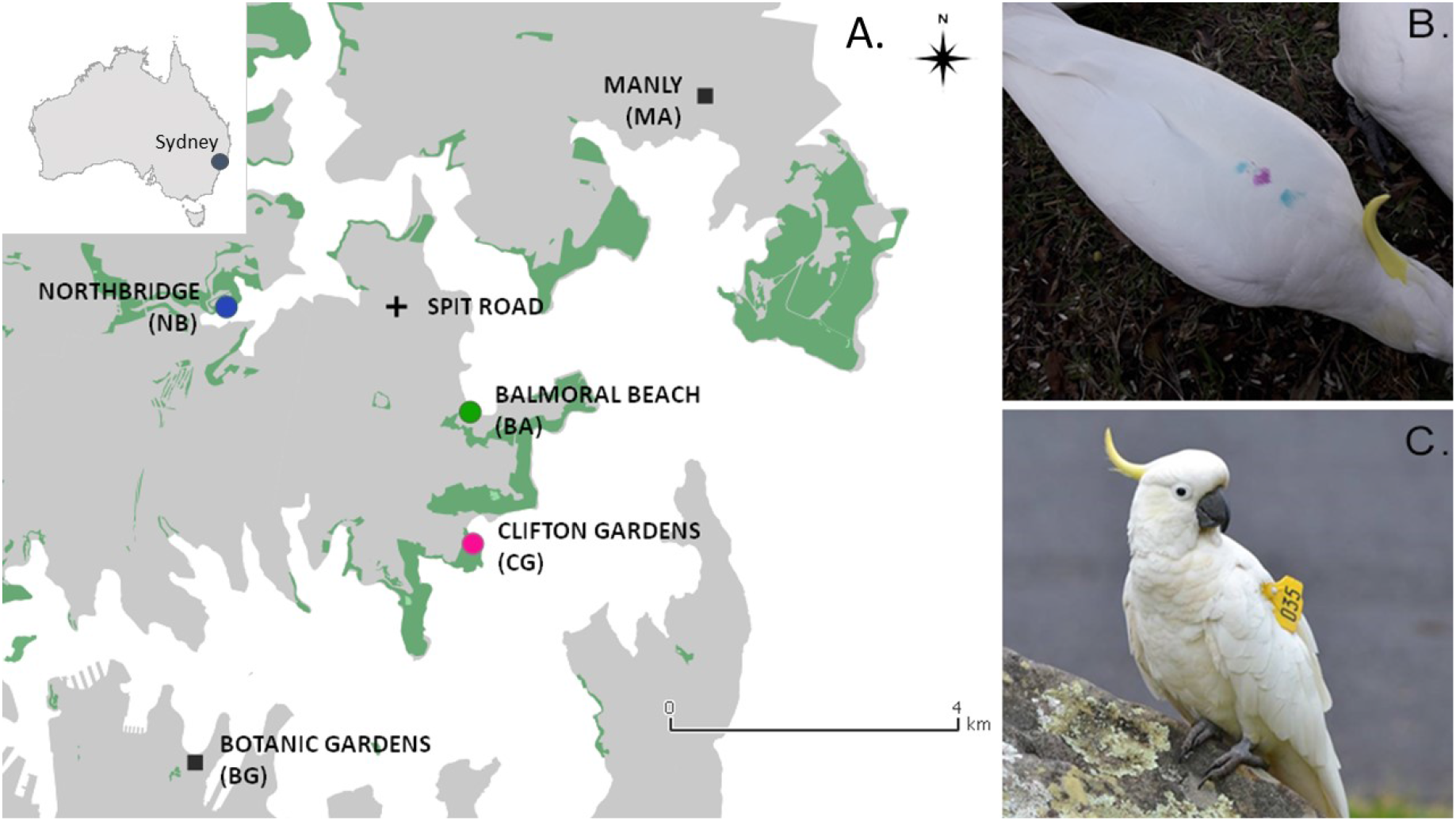
Study locations and examples of markings. A) Map of sampling locations. Water surfaces are represented in white, hard surfaces (e.g. buildings, roads) in grey, and vegetation (e.g. parks, bush-land) in green. Coloured circles represent focal roosts (green: BA, pink: CG, blue: NB) where social observations were conducted; grey squares are nearest neighbouring roosts (MA, BG) where birds were also marked and additional genetic samples were taken. The black cross represents an intermediate foraging site where birds from the three focal roosts mixed on a regular basis while being fed by locals, and social data was also collected. The map was constructed in QGIS (QGIS Development Team, 2009), using OpenStreetMap (OpenStreetMap contributors, 2017). B) Example of marking with temporary dye, “Teal-Pink-Teal”, the marks remain visible for approximately 3 months, C) Example of a wing-tagged individual 035, named “Shakespeare”.

Individuals were opportunistically sexed and aged by eye colour (brown = juvenile <7 years; red = adult female; black = adult male). Finally, feather samples were collected without capture by plucking feathers from the back of habituated individuals. Once collected, feather samples (n = 1-10 feathers per individual) were stored in paper envelopes until ready for processing. The feather tip (1-2mm; n = 351) was separated for genetic analysis (see below). This produced successful data for 244 samples; these were used to confirm morphometric sexing for adults, to assign sex for juveniles, and for relatedness analyses. All procedures were approved by the ACEC (ACEC Project No. 19/2107).

### Measuring social associations and interactions

We collected intensive social data at the three focal roost sites in northern Sydney and at one intermediate foraging site (Figure 1A) for ten days in July (July 8 - July 20) and ten days in September (September 19 - October 2) 2019. As for marking, individuals were attracted to the ground using small quantities of sunflower seeds scattered over a large area of approximately 385-500*m*^2^ to encourage ground foraging (e.g. of grass roots and shoots). Once the individuals were foraging on natural food sources, presence/absence scans (Altmann, 1974) were conducted every 10 minutes, where we recorded the identities of all individuals present within this study area in the park, both on the ground and in surrounding small trees.

Data was collected at the focal roosts over 3 hours (July 2019) or 2.5 hours (September 2019 - time at each group adjusted to fit within daylight) each day. Time periods for presence scans at the intermediate foraging site were more opportunistic, depending on the feeding regime of the local resident (no additional feeding or interference was undertaken). Altogether, the final dataset consisted of 819 scans.

In addition to social association data, affiliative and aggressive interactions were recorded *ad libitum* between scans (Altmann, 1974). The number for each type of interaction is presented in Table S2. Finally, we censused roost co-membership on three separate occasions (July 17th-21st, September 15th-20th, September 30th-October 3rd 2019) at the three focal roosts. The observer stood under the roost trees pre-dawn, and attracted individuals down to the ground with sunflower seed as soon as the birds began to become active, recording all marked individuals present. Roost counts were solely used to assign birds to a particular roost. Assignments were made if birds had been observed sleeping at one roost site at least twice (n=153 individuals for whom roost-memberships could be classified). Across the three focal roosts, marking location corresponded to roost-membership for 125 of 153 birds (Pearson’s Chi-squared test, *X*^2^=178.67, df=4, p<0.001).Marking location was therefore used as proxy for roost-membership for birds where roost was not known.

### Genetic Analysis

#### DNA analyses

DNA was processed by Diversity Arrays Technology (DArT), using their proprietary restriction-enzyme-based genome complexity reduction platform called DArTseq™ (Kilian et al., 2012; Cruz et al., 2013). Four combinations of restriction enzymes were tested and the PstI-SphI combination was selected. After digestion, DNA was processed following Kilian et al. (2012), using two different adaptors corresponding to two different restriction enzyme overhangs. Both adaptors contained an Illumina flow cell attachment sequence, and the PstI-compatible adapter also contained a sequencing primer sequence and a barcode region. A PCR was performed on the resultant libraries as follows: initial denaturation at 94°C for 1 min, then 30 cycles of 94°C for 20 sec, 58°C for 30 sec and 72°C for 45 sec, and a final extension step at 72°C for 7 min. After PCR, equimolar amounts of the libraries underwent a c-Bot (Illumina) bridge PCR followed by single end sequencing for 77 cycles on an Illumina Hiseq2500. Additional details on this platform can be found in Kilian et al. (2012) and Cruz et al. (2013).

The resultant sequence data were processed using the proprietary DArT Pty Ltd analytical pipelines. Poor quality sequences were removed, and stringent selection criteria was applied to the barcode region to de-multiplex the sequence reads (minimum barcode Phred score was set at 30, while the minimum whole-read Phred score was set at 10). Sequences were then truncated to 69 bp and clustered with a Hamming distance threshold of 3. Low quality bases from singleton tags were corrected based on collapsed tags. These sequences were used for SNP calling using the proprietary DArTsoft14 pipeline based on DArT PL’s C++ algorithm. True orthologous SNP variants were discriminated from paralogous sequences by assessing various parameters, including sequence depth, allele count and call rate. We calculated SNP error rates before and after filtering - i.e. the number of SNP mismatches between replicate pairs over the total number of SNPs that were not missing in both replicates. Analysis was based on the 13 replicate samples (Table S3) using R functions from Mastretta-Yanes et al. (2015).

#### SNP filtering

The DArTseq SNPs were filtered based on data quality, missing data and linkage using the R package dartR 1.0.5 (Gruber et al., 2018). The initial dataset comprised 58,740 SNPs. Loci that were genotyped in >80% of individuals were removed (i.e. call rate). DArT runs 30% of the samples in replicate in independent libraries and sequencing runs. Loci were removed that were not 100% reproducible, based on the consistency of the locus measured across these technical replicates. SNPs with a read depth <10 and a minor allele frequency <0.01 were removed. Putatively sex-linked markers based on SNPs that were mostly homozygous in males and mostly heterozygous in females, and/or SNPs that aligned with the Z-chromosome of the Finch genome were removed. Only one SNP was retained per DArTseq marker to remove potentially-linked SNPs. After implementing these filters, the dataset comprised 12,447 SNPs.

Initially, we did not filter for Hardy Weinberg Equilibrium (HWE) as the dataset likely contains numerous family groups, violating the HWE assumption of random mating. However, as a conservative approach, we subsequently re-analysed a dataset with HWE filtering implemented to check for consistency. We performed the HWE test using the R package pegas 0.13 (Paradis, 2010), implementing 1000 Monte Carlo replicates. Loci with a p-value <0.05 based on the exact p-test were removed (removing a further 3507 SNPs).

Finally, before further downstream analysis, one of each of the replicate samples (with the least amount of missing data) were removed from the dataset. We then excluded one potential hybrid individual and one individual with >60% missing data; all other individuals had <15% missing data. This resulted in 393 individuals for the relatedness analysis, of which 205 belonged to the 5 roosts considered in our study.

#### Relatedness analysis

We compared eight relatedness estimators to determine which one was the most appropriate for the SC-cockatoo SNP dataset (Wang, 2011). This included seven estimators implemented in the COANCESTRY software package (Wang, 2011), and a maximum likelihood kinship coefficient, calculated using the R package SNPRelate 1.14 (Zheng et al., 2012). To assess the suitability of these relatedness estimators, we simulated 100 parent-offspring pairs, 100 full-sibling pairs, 100 half-sibling pairs, 100 cousin pairs and 100 unrelated pairs and assessed the correlation between the relatedness estimates and true relatedness values. These simulations were performed in COANCESTRY using the empirical SNP allele frequencies and missing data. We compared the estimates using base R functions and the R package related 1.0 (Pew et al., 2015). After selecting the most reliable relatedness estimator based on the simulation data, we performed a relatedness analysis on the empirical dataset. The reliability of the estimator was verified by assessing how well it approximated two known parent-offspring relationships included in the dataset.

### Social network analyses

We considered four social networks: social associations, low- and high-intensity aggression networks, and allopreening. All social analyses were done using RStudio (version 1.4.1103; R version 4.0.4, R Core Team, 2021).

#### Social associations

To create social networks for foraging associations, we created a group by individual matrix, where individuals were assigned to the same group if observed in the same presence scan. Given that estimates for rarely observed individuals may be inaccurate (Farine and Whitehead, 2015), we thresholded our data following the example of Aplin et al. (2013). In our case, removing individuals with fewer than 22 observations meant there was no significant relationship between number of observations and the unweighted degree. We then used the simple ratio index (SRI; Whitehead, 2008) to create an undirected network for each time period (July & September 2019) where edges between individuals were scaled between 0 (never observed in the same group) and 1 (always observed together). The association matrices for each period were highly correlated (Mantel test, n=72, r=0.63, p<0.001). Therefore, association data for both study periods were pooled together for further analysis, resulting in a social network of 254 individuals. We tested for evidence of non-random preferred associations by examining whether the coefficient of variation (CV) of the observed SRI were significantly higher than permuted values. Permutations were conducted in the R package *asnipe* (Farine, 2013), permuting individuals in close spatio-temporal proximity, keeping rows and columns constant.

#### Aggression networks

To create aggression networks, we first *post hoc* split behavioural sequences of aggressive interactions into two distinct types of behaviours that clearly differed in risk: low intensity (no body contact, hereafter LI-) and high intensity (body contact & chases, hereafter HI-). One might expect different underlying decision rules to these interactions, especially in interactions with kin, as only high intensity interactions involve a risk of injury. For LI-aggressions this resulted in *N_int_*=1,457 interactions, and for HI-aggressions, this gave *N_int_*=91 interactions (Table S2). For each type of aggression, we then calculated an undirected interaction matrix that additionally controlled for social opportunities to interact. For the LI-aggressions, edge weights were obtained by calculating for each dyad the number of LI-aggressions divided by the total number of scans in which individuals were associated. For HI-aggressions, we calculated the proportion of HI-aggressions among all agonistic interactions between a given dyad, therefore controlling for the overall tendency of two individuals to engage in aggressive interactions.

#### Allopreening networks

Allopreening events were recorded *ad libitum* over the observation period (n=26). Within an allopreening event, preenings were mostly reciprocated. They also occurred in trees, making it challenging to record the initiator of the event. To create affiliative networks, we therefore constructed undirected networks where edge weights were obtained by calculating for each dyad the number of preening interactions divided by the sum of all preening events for both individuals. As observations of allopreening events were relatively rare (Table S2), we did not threshold the network.

### Statistical analysis of kinship and social networks

#### Genetic differentiation between roosting communities

To examine whether there was any genetic differentiation between roosts, we performed two analyses: the first included five roosting communities, located within a 10 kilometer radius (BA, CG, NB, BG, MA - Figure 1). The second focused on the three focal roosting communities (BA, CG, NB - Figure 1) located within a four kilometer radius. For each analysis, we used mantel tests in the R package *vegan* (Oksanen et al., 2013), to correlate relatedness matrices with a spatial matrix of roost membership. This was performed using roost co-membership assigned for individuals at the three focal roosts by roost counts.

#### Influence of kinship on social interactions

##### Level 1: study population

To test whether associations, LI- and HI-aggressive interactions and/or affiliative interactions were influenced by relatedness, we compared kinship with these measures in separate models using multiple regression quadratic assignment procedures (MRQAP) via the *asnipe* package (Farine, 2013), where the significance was ascertained using permutations (Farine and Carter, 2022). We additionally included roost co-membership and sex- and age-similarity as covariates in all models. In all MRQAP analyses, we used a continuous relatedness matrix. However, because we considered it unlikely for individuals to recognize second- or third-degree relatives, we also re-ran the same analysis on a binary matrix, specifying whether or not a given dyad were first degree relatives, i.e. that their relatedness was above or equal to 0.36 (Figure S1; Tables S4, S5, S6, S7).

##### Level 2: Within vs between roost interactions

To test whether the correlation between relatedness and our social measures was mostly driven by within-roost or between-roost interactions, we conducted a second analysis where we: (i) subset all edges to within-roost only and restricted randomisations to within the same roost, and (ii) subset all edges to those occurring between individuals of different roost membership. Due to an increased likelihood of type II errors with this reduced dataset, we did not do this analysis for HI-aggression or allopreening. For all analysis, an estimate of type II errors (calculated following Foroughirad et al., 2019) are presented in Section S1.

## Results

### Social organisation

In northern Sydney, 561 birds were individually marked or recognisable, and genetic material was collected from 205 birds. We sampled 164 birds from the three focal roost sites (Figure 1A; BA: n=68; CG: n=55; NB: n=41), comprising 63 females and 72 males, of which 83 were adults, 45 juveniles and 37 birds of unknown age (Figure 2). At two additional roost sites, we sampled 41 birds (BG: n=17; MA: n=24), comprising 12 females and 23 males; 18 adults, 17 juveniles, and 6 birds of unknown age. Of the eight relatedness estimators compared, the SNPRelate maximum likelihood kinship coefficient was most appropriate for this SNP dataset based on simulated dyads (Figure S1), having the highest correlation with the expected relatedness values (cor = 0.999; p<0.001).

**Figure 2:**
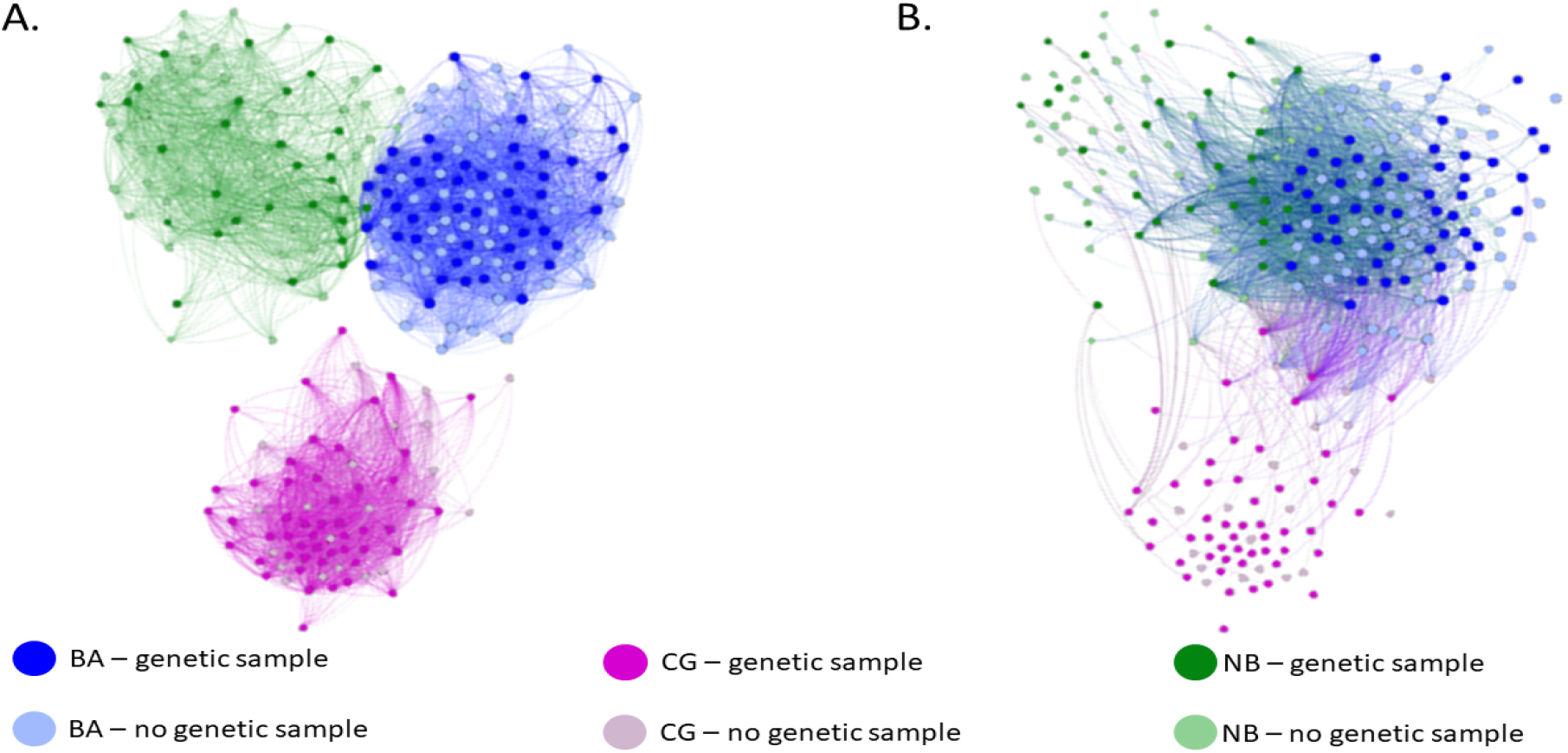
Association network of Sulphur-crested cockatoos across three roosting locations, with A) representing associations between individuals of the same roost, and B) representing only associations between individuals of different roosts. Networks are thresholded to only include individuals with at least 22 observations. Nodes are coloured by roost (blue: BA, pink: CG, green: NB), and the shade of each colour illustrates whether a genetic sample has been collected (dark shades) or not (lighter shades). Edges between nodes represent association strength, and are thresholded to above 0.15 for visual clarity.

Overall, there was a weak tendency for individuals to be more related within than between roosting sites, but this was only present when considering all 5 roosts (focal roosts: Mantel Test, n=164, r=−0.003, p=0.60; all five roosts: Mantel Test, n=205, r=0.011, p=0.049). The higher relatedness within - compared to between - roosts was not solely due to delayed dispersal in juveniles; within each roost site, the number of adult-adult close-kin dyads was higher than expected within each population under random mixing (t-test, t=2.28, df= 82, p=0.025). In addition, juveniles tended to be observed at more roosting sites than adults (Figure S2A; females estimate ± 95% confidence interval: ES= 0.553 [0.063-1.04], p=0.02, males: ES=0.739 [0.256-1.222], p<0.001; p-values calculated using Tukey adjustments for multiple testing). Juvenile females, but not juvenile males, tended to be less faithful to a specific site than adults of the respective sex, where faithfulness was defined as the proportion of observations at an individuals assigned roost (Figure S2B; females: ES=−0.107 [−0.216-0.002], p=0.06, males: ES=−0.058 [−0.151-0.035], p=0.37, p-values calculated using Tukey adjustments for multiple testing).

Analysis of association patterns revealed strong evidence for non-random associations (Median CV random = 1.26, CV obs=1.25, P=0.003), suggesting that individuals had preferred and non-preferred associates. While the social association network was clearly clustered by roosting community (Figure 2A), individuals also exhibited extensive foraging associations with individuals from different roost groups (Figure 2B).

### Relatedness and social networks

When the entire study population was considered, kinship positively predicted social association strength (Figure 2A & B, ES=0.097, p<0.001). As expected, associations were also stronger between members of the same roost and with individuals of similar age (Figure 2, roost co-membership: ES=0.097, p<0.001, age: ES=0.006, p=0.012), but not of the opposite sex (sex: ES=0.002, p=0.18). The results were similar when considering only within-roost edges (kinship: ES=0.11, p=0.005; sex: ES=0.006, p=0.24; age: ES=0.018, p<0.001; Figures 2A, 3A). When sub-setting the dataset to edges between individuals of different roost locations (Figure 2B), the effect of kinship on association strength was still present (ES=0.086, p=0.001), but there with no effect of sex or age (sex: ES=−0.001, p=0.58; age: ES=0.0003, p=0.87). Finally, results were similar whether kinship was considered as a continuous or binary measure (close-kin or not) (Table S4).

### Relatedness and aggressive interactions

Even after accounting for social opportunity, kinship was positively correlated with LI-aggression (ES=0.41, p=0.020, Figure 3B). LI-aggressions (*N_int_*=1,457, Table S2) also occurred more frequently between individuals belonging to the same roost than with individuals visiting from different roosts (roost co-membership: ES=0.17, p<0.001). There was also a trend for age similarity to predict LI-aggressions (ES=0.020, p=0.09), but no effect of sex (ES=0.01, p=0.23). When considering only within-roost edges, the results were similar (kinship: ES=0.78, p=0.03; age: ES=0.063, p=0.08; sex: ES=0.038, p=0.31), as were the results when only considering close-kin dyads (Table S5). Between-roost LI-aggressions, however, were not influenced by kinship, age or sex (kinship: ES=−0.025, P=0.41; age: ES=0.002, p=0.72; sex: ES=0.004, p=0.41), though there was a tendency to avoid directing LI-aggressions towards close-kin of different roost-membership (ES=−0.010, p=0.08, Table S5).

**Figure 3:**
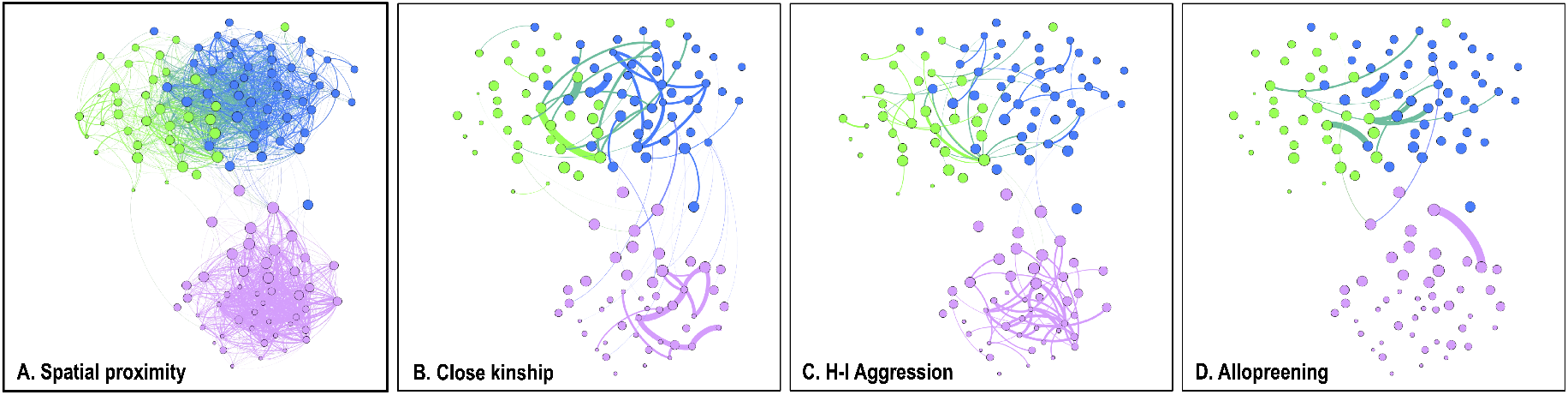
Social networks across three roosting communities. A) Spatial association network of cockatoos where edge width represents association strength. Edges are thresholded to above 0.15 for visual clarity. B) Association network subset to show edges between individuals with close kinship. C) Association network subset to show edges between individuals that engaged in HI-aggressions. D) Association network subset to show edges between individuals that exchanged in allopreening. In all networks, node size represents average association strength, nodes are coloured by roost (blue: BA, pink: CG, green: NB), and networks are thresholded to only include individuals with at least 22 observations.

Similar to LI-aggressions, HI-aggressive interactions (*N_int_*=91, Table S2) occurred more frequently between individuals belonging to the same roost, rather than between residents and individuals visiting from different roosts (roost co-membership: ES=0.01, p<0.001). Kinship (ES=0.055, p=0.03) also positively predicted HI-aggression (Figure 3C), but there was no clear effect of age or sex similarity (age: ES=0.003, P=0.17; sex: ES=0.003, P=0.09). Due to low sample size, we did not examine the interaction between kinship and roost membership on HI-aggressions.

### Relatedness and allopreening interactions

Observations of allopreening events were scarce, with 26 allopreening interactions involving 41 individuals (Table S2). Yet even so, individuals were observed engaging in allopreening events with multiple partners. Allopreening interactions were more likely to be observed between stronger social associates (ES=0.191, p<0.001, Figure 3), related individuals (ES=0.275; p=0.008), individuals of different roost-membership (ES=−0.015, p=0.02), and tended to be more frequent between members of the opposite sex (ES=0.012, p=0.08).

## Discussion

By combining relatedness data with extensive observations of social networks and interactions at multiple levels, our study gained an unprecedented insight into effects of kinship on the social lives of a wild parrot species. First, we collected genetic material and built kinship networks for five neighbouring roosting communities in a 25km^2^ urban and suburban area of Sydney. Our results showed a very weak genetic differentiation between these roost sites, suggesting extensive gene flow. This was supported by our repeated observations of between-roost movements, especially for juvenile females. Second, we used social network analysis to show that despite well-mixed social networks and a low degree of kin structuring, individuals preferentially associated with close kin. Supporting this, individuals were also more likely to allopreen kin, an important affiliative behaviour in parrots (Morales Picard et al., 2020). Contrary to expectations, aggressive interactions were also more likely to occur between kin within roost sites; intriguingly this was absent from between-roost interactions. Finally, we showed that this preference to associate with kin could be observed both within roost sites, and in encounters with individuals from other roosting sites.

A previous study in this system has shown that individuals maintain long-term preferred social associations (Aplin et al., 2021). Our work extends those findings to reveal one driver of these association choices, with kinship predicting social associations within roost sites, suggesting an active preference to associate with kin. Perhaps even more interestingly, we found that kinship was also positively correlated with association strength between individuals of different roost groups, suggesting that individuals continued to recognise and prefer kin even after dispersal. One benefit of such kin-based between-roost associations may be facilitated emigration into, or interactions with, groups containing closely related individuals (Van Hoof, 2000). For example, in cooperatively breeding brown jays (*Cyanocorax morio*), males tend to disperse into groups where male relatives are already established (Williams and Rabenold, 2005). In the case of the brown jay, the formation of alliances between related males may facilitate the entrance into the new social group. A similar behaviour was also described for invasive monk parakeets, where male siblings exhibited parallel dispersal, then often cooperated when building a new nest and breed (Dawson Pell et al., 2021). Cooperative breeding or the formation of alliances has not been observed in SC-cockatoos, yet the observed preening interactions both between kin and individuals of different roosts is an intriguing indicator of the potential importance of maintaining relationships across groups.

In birds, preening is often described as having an essential role in maintaining pair bonds, such as mate bonding and partner retention across breeding seasons (Gill, 2012), or parental cooperation (Kenny et al., 2017). While our study does indeed identify allopreening between members of different sexes, it was not restricted to putative mates. Rather, individuals were observed engaging in preening events with multiple partners, and these interactions were positively with both kinship and social association strength. Allopreening is generally understudied in wild parrots (but see McCulloch, 1981; Diamond and Bond, 1999; Taylor, 2002). In captivity, allopreening patterns of nine parrot species were positively correlated to the time a dyad spent in close proximity (Morales Picard et al., 2020). Furthermore, a preference to preen related, rather than unrelated individuals, has been described in juvenile birds (e.g. corvids: Fraser and Bugnyar, 2010; e.g. captive parrots: Garnetzke-Stollmann and Franck, 1991), where it is hypothesised to help build bonds with important social partners. The number of preening events in our dataset was low, and conclusions need to be drawn with care. Yet our results suggest that SC-cockatoos may gain some direct or indirect benefits from building and maintaining strong social bonds with kin, even after dispersal. Future work should seek to quantify what benefit these between-roost kin relationships give to individuals.

Somewhat surprisingly, we found that SC-cockatoos directed aggression towards individuals within rather than outside their roost. Furthermore, after social associations were controlled for, individuals also preferred to aggress kin with which they shared roost co-membership. A preference to direct aggression towards familiar individuals could potentially reflect strategies for improving dominance (Hobson et al., 2014; Hobson et al., 2021), especially since 93% of all observed aggressive interactions involved only displacements that bear no direct risks of injuries for the individuals involved. However the increased aggression between kin was in complete contrast to our prediction of higher tolerance. To note, all juveniles in our dataset were already independent of adults (2 to 7 years of age). One potential explanation therefore might come from the aggressive eviction hypothesis (Marsh, 1988, Wahlström, 1994), which states that increased aggression can act to drive emigration. For example, in western bluebirds (*Sialia mexicana*), dispersal, and dispersal distance, is influenced by aggressions from kin (Aguillon and Duckworth, 2015). Although this hypothesis was formulated for natal emigration, it seems reasonable to expect increased intra-group aggression to also drive secondary dispersals, and even short-term movements between social groups. Unfortunately we did not have a large enough sample size to classify and distinguish between aggressive interactions between offspring-parents and between siblings. However this hypothesis would be consistent with our observation that this preference to aggress kin is either lost, or perhaps reversed, when individuals interact with those of different roost-membership.

Overall, the social system of SC-cockatoos allowed us to disentangle association preferences from spatial overlap, with well-mixed social networks within clearly-defined roosting groups coupled with regular associations with individuals of different roosts. We found that, despite exhibiting low overall kin structuring, individuals prefer to associate and allopreen with their kin both within and between roosting groups. This therefore suggests that: 1) individuals are able to recognise kin, including siblings, and actively prefer them, and 2) that kin-recognition is retained over a considerable period of time, including after dispersal. In passerine birds, cross-fostering experiments have demonstrated that kin recognition is often learned in the nest (Sharp et al., 2005). Yet the typical clutch size for SC-cockatoos is one or two chicks per year, and juveniles appear to be largely independent at one year of age, suggesting that kin recognition must also occur across broods. In zebra finches between-brood recognition is achieved through olfactory cues (Krause et al., 2012). However, given the evidence for extensive vocal learning in parrots (Wright and Dahlin, 2018), and the existence of parent-facilitated vocal signatures in parrotlets (*Forpus passerinus*, Berg et al., 2012; Berg et al., 2013), it seems most likely that kin recognition in SC-cockatoos involves vocal learning.

Whatever the mechanism of kin recognition, preferred associations with kin are often thought to benefit individuals in a number of ways, including through inclusive fitness, increased tolerance and cooperation. What the benefits are in this case are not obvious – kin do not cooperate to breed, and rather than exhibiting increased tolerance, individuals actually appear to be more likely to aggress kin, including with high intensity aggressions that can carry risk of injury. While we were unable to ascertain the drivers, our study did reveal that interactions with related individuals are context-dependent, suggesting the existence of interaction rules that vary with the behavioural context and on roost membership. More investigation of these dependencies may therefore be a fruitful avenue to identify the benefits of kin relationships in this system. Perhaps most intriguingly, our results suggest that roost membership is a key factor, with SC-cockatoos exhibiting preferred associations with individuals of other roosting groups that are often kin based, and appearing to maintain these relationships with affiliative behaviours such as allopreening. How individuals benefit from these extended networks remains to be further explored.

## Acknowledgements

We would like to acknowledge the Gamaragal and Gadigal people of the Eora Nation as the Traditional Custodians of the Land on which this study was conducted.. We pay our respects to the Elders past, present, and emerging.

Thank you to Damien Farine for discussions on analysis. Thank you to Eliane McCarthy for assistance in preparing the genetic samples. Finally, we would like to thank the community of citizen scientists participating in the Wingtags and Big City Birds project. Last, but not least, we would like to thank the two anonymous reviewers who’s thoughtful comments greatly greatly helped improving the manuscript.

## Authorship contributions

J.P., L.M.A., J.M.M. conceived the ideas and designed the study; J.P., L.M.A., B.C.K. and J.M.M. collected social and morphometric data and collected feather samples. K.M.E collated and analysed genetic data, and J.P. and L.M.A. performed analysis on sociality and kinship. All authors contributed to writing the manuscript and gave final approval for publication.

## Data Availability Statement

Upon acceptance, the data supporting this manuscript will be made available at Edmond, the open access repository of the Max Planck Society, and the DOI provided in the manuscript.

## Supplementary Materials

### S1 Sensitivity analysis

To assess the sensitivity of our analysis, we assessed the type II error rates following (Foroughirad et al., 2019). We run 1,000 iterations of each MRQAP in the original analysis. For each iteration, we extracted the significance of kinship on the response variable, using MRAQP (significance calculated on 1,000 permutations following Farine and Carter, 2022). We then calculated the type II error rates, where we considered p-values above alpha=0.05 to result in a type II error when the original result reported in the paper was significant.

**Table S1:** Estimated type II error rates for each analysis presented in section. Error rates between parentheses represent type II error rates in cases where the original result (reported in section) was non-significant, and vice-versa.

| type | type II error (%) |
| --- | --- |
| association |  |
| *within roosts* | 0.1 |
| *between roosts* | 1 |
| LI-aggression |  |
| *within roosts* | 6.6 |
| *between roosts* | (0) |
| HI-aggression |  |
| *study population (no subset)* | 6.7 |
| Preening |  |
| *study population (no subset)* | 5.1 |

### S2 Figures and tables

**Figure S1:**
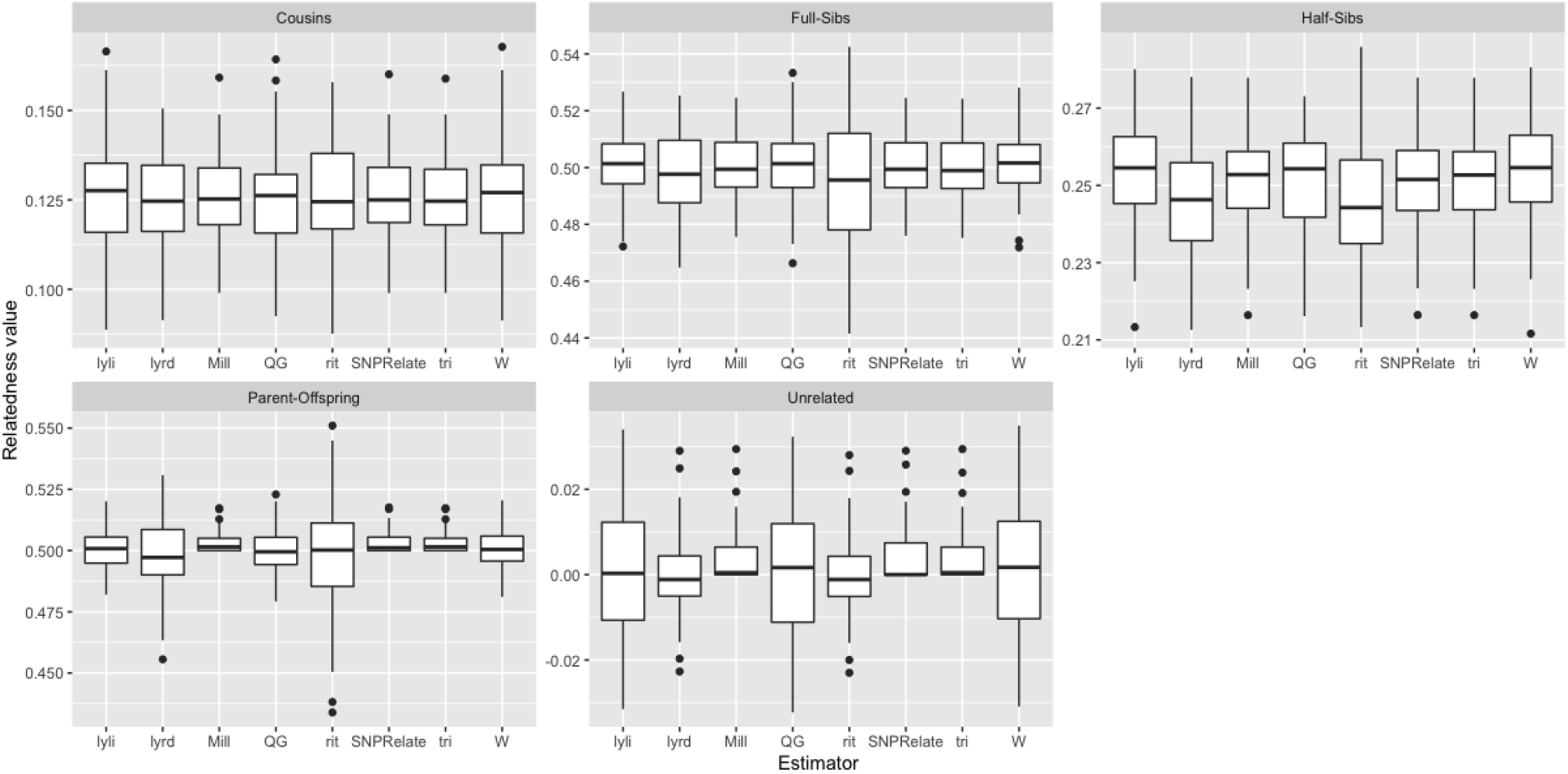
Distribution of relatedness values for eight relatedness estimators based on simulated data of five relationship types. lyli = Li et al. estimator; lyrd = Lynch and Ritland estimator; Mill = dyadic likelihood estimator; QG = Queller and Goodnight estimator; Rit = Ritland estimator; SNPRelate = SNPRelate maximum likelihood kinship coefficient; tri = Triodic likelihood estimator; W = Wang estimator.

**Table S2:** Summary of the social data The number of observations (namely number of scans for the associations, number of interactions for preening and aggression) and number of individuals (with sex and age: J=juvenile, A=adults, U=unknown age) involved for each behaviour analysed in this study. Only individuals with at least 22 sightings are included.

| Level of Analysis | Behaviour | n females<br>(J/A/U) | n males (J/A/U) | n obs |
| --- | --- | --- | --- | --- |
| Population | Association | 73 (20/35/2) | 72 (25/37/10) | 864 |
| Population | Preening | 16 (6/7/3) | 25 (7/10/8) | 26 |
| Within- & between-roost | Aggression (low intensity) | 56 (18/28/10) | 74 (21/38/15) | 1,457 |
| Population | Aggression (high intensity) | 56 (18/28/10) | 74 (21/38/15) | 91 |

**Figure S2:**
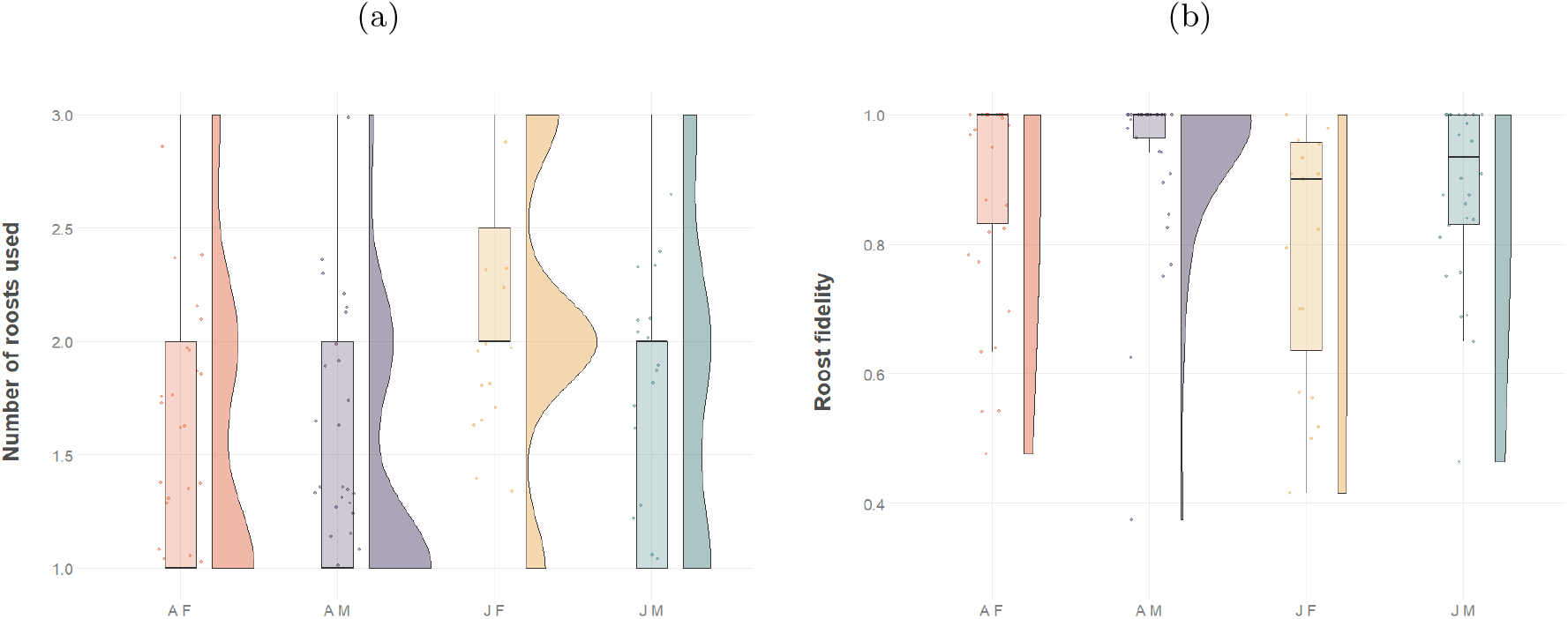
Between roost movements of SC-cockatoos according to their age-sex category. AF: adult females, AM: adult males, JF: juvenile females, JM: juvenile males. (a) Number of roosts used by individuals of each age sex group (maximum 3 roosts). (b) roost preference for each age-sex group. Preference is defined as the proportion of observations for a focal individual at its assigned roost by roost count, over to the total number of observations of this individual during foraging associations.

**Table S3:** Allele error rate calculated as the total number of alleles that mismatch between the replicate pair divided by the total number loci that were not missing in both samples. NB: some replicates have the same names, while others have different names, and triplicates include three pairwise comparisons. The high error rate in two of the S111 pairwise comparisons suggests a mislabelling issue for one sample, or a failed assay.

| Replicate name | Allele error rate for full dataset (58,740 SNP) | Allele error rate for filtered dataset (12,447 SNPs) |
| --- | --- | --- |
| S111 triplicate 1 | 0.200 | 0.282 |
| S111 triplicate 2 | 0.023 | 0.010 |
| S111 triplicate 3 | 0.200 | 0.289 |
| S42/S160 replicate | 0.018 | 0.011 |
| S149 replicate | 0.012 | 0.007 |
| *tag<sub>0</sub>25a/S281replicate* | 0.012 | 0.003 |
| *tag<sub>1</sub>07replicate* | 0.009 | 0.002 |
| *tag<sub>1</sub>07/S254replicate1* | 0.032 | 0.032 |
| *tag<sub>1</sub>07/S254replicate2* | 0.033 | 0.031 |
| S379 replicate | 0.017 | 0.008 |
| *tag<sub>0</sub>35/S345replicate* | 0.011 | 0.002 |
| S295 replicate | 0.023 | 0.010 |
| S302/S344 replicate | 0.017 | 0.003 |

**Table S4:** Within and between roost associations predicted by close-kinship, sex and age.

| type | coefficients | estimate | pvalue |
| --- | --- | --- | --- |
| overall | intercept | 0.028 | <0.001 |
|  | kin (binary) | 0.052 | <0.001 |
|  | sex | 0.003 | 0.10 |
|  | age | 0.006 | 0.01 |
|  | roost | 0.097 | 0.01 |
| between roosts | intercept | 0.031 | <0.001 |
|  | kin (binary) | 0.022 | 0.04 |
|  | sex | / | / |
|  | age | / | / |
| within roots | intercept | 0.118 | 0.001 |
|  | kin (binary) | 0.037 | 0.04 |
|  | sex | / | / |
|  | age | / | / |

**Table S5:** Low-intensity aggressions at roost level, predicted by close-kinship, roost-membership, age and sex.

| type | coefficients | estimate | pvalue |
| --- | --- | --- | --- |
| overall | intercept | -0.008 | 0.42 |
|  | kin (binary) | 0.095 | 0.09 |
|  | sex | 0.016 | 0.15 |
|  | age | 0.020 | 0.091 |
|  | roost | 0.16 | 0.001 |
| within roost | intercept | 0.13 | <0.001 |
|  | kin (binary) | 0.10 | 0.08 |
|  | sex | / | / |
|  | age | / | / |

**Table S6:** High-intensity aggressions at roost level, predicted by close-kinship measures, roost-membership, age and sex.

| type | coefficients | estimate | pvalue |
| --- | --- | --- | --- |
| overall | intercept | -0.0002 | 0.97 |
|  | kin (binary) | 0.040 | 0.04 |
|  | sex | 0.004 | 0.06 |
|  | age | 0.003 | 0.17 |
|  | roost | 0.011 | 0.001 |

**Table S7:**
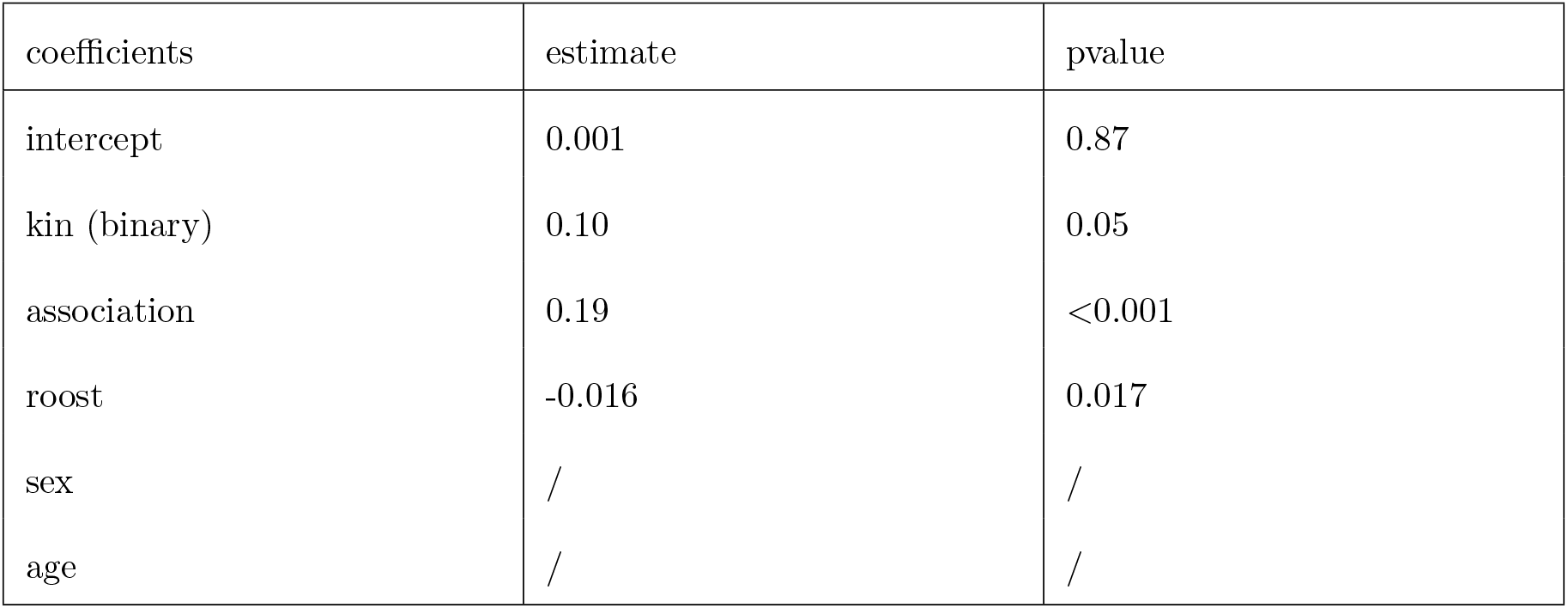
Allopreening interactions predicted by close-kinship, associations, roost-membership, age and sex.

## References

Aguillon, S. M., & Duckworth, R. A. (2015). Kin aggression and resource availability influence phenotype-dependent dispersal in a passerine bird. Behavioral Ecology and Sociobiology, 69 (4), 625–633.

Altmann, J. (1974). Observational study of behavior: Sampling methods [Publisher: Brill Section: Behaviour]. Behaviour, 49 (3), 227–266. https://doi.org/10.1163/156853974X00534

Aplin, L. M., Farine, D. R., Morand-Ferron, J., Cole, E. F., Cockburn, A., & Sheldon, B. C. (2013). Individual personalities predict social behaviour in wild networks of great tits (parus major). Ecology letters, 16 (11), 1365–1372.

Aplin, L. M., Major, R. E., Davis, A., & Martin, J. M. (2021). A citizen science approach reveals long-term social network structure in an urban parrot, cacatua galerita. Journal of Animal Ecology, 90 (1), 222–232.

Berg, K. S., Beissinger, S. R., & Bradbury, J. W. (2013). Factors shaping the ontogeny of vocal signals in a wild parrot. Journal of Experimental Biology, 216 (2), 338–345.

Berg, K. S., Delgado, S., Cortopassi, K. A., Beissinger, S. R., & Bradbury, J. W. (2012). Vertical transmission of learned signatures in a wild parrot. Proceedings of the Royal Society B: Biological Sciences, 279 (1728), 585–591.

Birch, J. (2017). The inclusive fitness controversy: Finding a way forward. Royal Society Open Science, 4 (7), 170335.

Blanco, G., Morinha, F., Roques, S., Hiraldo, F., Rojas, A., & Tella, J. L. (2021). Fine-scale genetic structure in the critically endangered red-fronted macaw in the absence of geographic and ecological barriers. Scientific reports, 11 (1), 1–17.

Brown, C. R., & Brown, M. B. (1987). Group-living in cliff swallows as an advantage in avoiding predators. Behavioral Ecology and Sociobiology, 21 (2), 97–107.

Carter, K. D., Seddon, J. M., Frère, C. H., Carter, J. K., & Goldizen, A. W. (2013). Fission–fusion dynamics in wild giraffes may be driven by kinship, spatial overlap and individual social preferences. Animal Behaviour, 85 (2), 385–394.

Chiyo, P. I., Archie, E. A., Hollister-Smith, J. A., Lee, P. C., Poole, J. H., Moss, C. J., & Alberts, S. C. (2011). Association patterns of african elephants in all-male groups: The role of age and genetic relatedness. Animal Behaviour, 81 (6), 1093–1099.

Croft, D., James, R., Ward, A., Botham, M., Mawdsley, D., & Krause, J. (2005). Assortative interactions and social networks in fish. Oecologia, 143 (2), 211–219.

Cruz, V. M. V., Kilian, A., & Dierig, D. A. (2013). Development of dart marker platforms and genetic diversity assessment of the us collection of the new oilseed crop lesquerella and related species. PloS one, 8 (5), e64062.

Dadda, M., Pilastro, A., & Bisazza, A. (2005). Male sexual harassment and female schooling behaviour in the eastern mosquitofish. Animal Behaviour, 70 (2), 463–471. https://doi.org/10.1016/j.anbehav.2004.12.010

Davis, A., Major, R. E., Taylor, C. E., & Martin, J. M. (2017). Novel tracking and reporting methods for studying large birds in urban landscapes [Publisher: Nordic Board for Wildlife Research]. Wildlife Biology, 2017 (4).

Dawson Pell, F. S., Senar, J. C., Franks, D. W., & Hatchwell, B. J. (2021). Fine-scale genetic structure reflects limited and coordinated dispersal in the colonial monk parakeet, myiopsitta monachus. Molecular Ecology, 30 (6), 1531–1544.

Diamond, J., & Bond, A. B. (1999). Kea, bird of paradox. University of California Press.

Evans, J. C., Lindholm, A. K., & König, B. (2021). Long-term overlap of social and genetic structure in free-ranging house mice reveals dynamic seasonal and group size effects. Current zoology, 67 (1), 59–69.

Farine, D. R. (2013). Animal social network inference and permutations for ecologists in r using asnipe. Methods in Ecology and Evolution, 4 (12), 1187–1194.

Farine, D. R., & Carter, G. G. (2022). Permutation tests for hypothesis testing with animal social network data: Problems and potential solutions. Methods in Ecology and Evolution, 13 (1), 144–156.

Farine, D. R., & Whitehead, H. (2015). Constructing, conducting and interpreting animal social network analysis. Journal of animal ecology, 84 (5), 1144–1163.

Foroughirad, V., Levengood, A. L., Mann, J., & Frère, C. H. (2019). Quality and quantity of genetic relatedness data affect the analysis of social structure. Molecular ecology resources, 19 (5), 1181–1194.

Fraser, O. N., & Bugnyar, T. (2010). The quality of social relationships in ravens. Animal behaviour, 79 (4), 927–933.

Garnetzke-Stollmann, K., & Franck, D. (1991). Socialisation tactics of the spectacled parrotlet (forpus conspicillatus). Behaviour, 119 (1-2), 1–29.

Gerlach, G., Hodgins-Davis, A., MacDonald, B., & Hannah, R. C. (2007). Benefits of kin association: Related and familiar zebrafish larvae (danio rerio) show improved growth. Behavioral Ecology and Sociobiology, 61 (11), 1765–1770. https://doi.org/10.1007/s00265-007-0409-z

Gill, S. A. (2012). Strategic use of allopreening in family-living wrens. Behavioral Ecology and Sociobiology, 66 (5), 757–763.

Grafen, A. (1984). Natural selection, kin selection and group selection [polistes fuscatus, wasps]. Behavioural ecology: an evolutionary approach/edited by JR Krebs and NB Davies.

Gruber, B., Unmack, P. J., Berry, O. F., & Georges, A. (2018). Dartr: An r package to facilitate analysis of snp data generated from reduced representation genome sequencing. Molecular Ecology Resources, 18 (3), 691–699.

Hamilton, W. D. (1964). The genetical evolution of social behaviour. ii. Journal of theoretical biology, 7 (1), 17–52.

Hardy, J. W. (1965). Flock social behavior of the orange-fronted parakeet. The Condor, 67 (2), 140–156.

Hepper, P. G. (1986). Kin recognition: Functions and mechanisms a review. Biological Reviews, 61 (1), 63–93.

Hobson, E. A., Avery, M. L., & Wright, T. F. (2014). The socioecology of monk parakeets: Insights into parrot social complexity. The Auk, 131 (4), 756–775. https://doi.org/10.1642/AUK-14-14.1

Hobson, E. A., Mønster, D., & DeDeo, S. (2021). Aggression heuristics underlie animal dominance hierarchies and provide evidence of group-level social information [Publisher: National Academy of Sciences Section: Biological Sciences]. Proceedings of the National Academy of Sciences, 118 (10). https://doi.org/10.1073/pnas.2022912118

Ilany, A., Holekamp, K. E., & Akçay, E. (2021). Rank-dependent social inheritance determines social network structure in spotted hyenas. Science, 373 (6552), 348–352.

Kenny, E., Birkhead, T. R., & Green, J. P. (2017). Allopreening in birds is associated with parental cooperation over offspring care and stable pair bonds across years. Behavioral Ecology, 28 (4), 1142–1148.

Kilian, A., Wenzl, P., Huttner, E., Carling, J., Xia, L., Blois, H., Caig, V., Heller-Uszynska, K., Jaccoud, D., Hopper, C., et al. (2012). Diversity arrays technology: A generic genome profiling technology on open platforms. Data production and analysis in population genomics (pp. 67–89). Springer.

Kirksey, E., Munro, P., van Dooren, T., Emery, D., Maree Kreller, A., Kwok, J., Lau, K., Miller, M., Morris, K., Newson, S., Olejniczak, E., Ow, A., Tuckson, K., Sannen, S., & Martin, J. (2018). Feeding the flock: Wild cockatoos and their facebook friends [Publisher: SAGE Publications Ltd STM]. Environment and Planning E: Nature and Space, 1 (4), 602–620. https://doi.org/10.1177/2514848618799294

Klauke, N., Schaefer, H. M., Bauer, M., & Segelbacher, G. (2016). Limited dispersal and significant fine - scale genetic structure in a tropical montane parrot species [Publisher: Public Library of Science]. PLOS ONE, 11 (12), e0169165. https://doi.org/10.1371/journal.pone.0169165

Klump, B. C., Martin, J. M., Wild, S., Hörsch, J. K., Major, R. E., & Aplin, L. M. (2021). Innovation and geographic spread of a complex foraging culture in an urban parrot. Science, 373 (6553), 456–460.

Krause, J., Krüger, O., Kohlmeier, P., & Caspers, B. A. (2012). Olfactory kin recognition in a songbird. Biology letters, 8 (3), 327–329.

Krause, J., Ruxton, G. D., Ruxton, G., Ruxton, I. G., et al. (2002). Living in groups. Oxford University Press.

Krützen, M., Sherwin, W. B., Connor, R. C., Barré, L. M., Van de Casteele, T., Mann, J., & Brooks, R. (2003). Contrasting relatedness patterns in bottlenose dolphins (tursiops sp.) with different alliance strategies. Proceedings of the Royal Society of London. Series B: Biological Sciences, 270 (1514), 497–502.

Kurvers, R. H., Adamczyk, V. M., Kraus, R. H., Hoffman, J. I., Van Wieren, S. E., Van Der Jeugd, H. P., Amos, W., Prins, H. H., & Jonker, R. M. (2013). Contrasting context dependence of familiarity and kinship in animal social networks. Animal Behaviour, 86 (5), 993–1001.

Levin, S. R., & Grafen, A. (2019). Inclusive fitness is an indispensable approximation for understanding organismal design. Evolution, 73 (6), 1066–1076.

Lindenmayer, D. B., Pope, M., Cunningham, R. B., Donnelly, C. F., & Nix, H. A. (1996). Roosting of the sulphur-crested cockatoo cacatua galerita. Emu-Austral Ornithology, 96 (3), 209–212.

Madden, J. R., Nielsen, J. F., & Clutton-Brock, T. H. (2012). Do networks of social interactions reflect patterns of kinship? Current Zoology, 58 (2), 319–328.

Marsh, C. (1988). Bb smuts, dl cheney, rm seyfarth, rw wrangham & tt struhsaker (eds) 1987. primate societies. chicago university press. Journal of Tropical Ecology, 4 (3), 317–318.

Mastretta-Yanes, A., Arrigo, N., Alvarez, N., Jorgensen, T. H., Piñero, D., & Emerson, B. C. (2015). Restriction site-associated dna sequencing, genotyping error estimation and de novo assembly optimization for population genetic inference. Molecular ecology resources, 15 (1), 28–41.

McCulloch, E. M. (1981). Short notes-mutual preening by four musk lorikeets. Australian Bird Watcher, 9 (4), 135.

Metheny, J. D., Kalcounis-Rueppell, M. C., Willis, C. K., Kolar, K. A., & Brigham, R. M. (2008). Genetic relationships between roost-mates in a fission–fusion society of tree-roosting big brown bats (eptesicus fuscus). Behavioral Ecology and Sociobiology, 62 (7), 1043–1051.

Morales Picard, A., Mundry, R., Auersperg, A. M., Boeving, E. R., Boucherie, P. H., Bugnyar, T., Dufour, V., Emery, N. J., Federspiel, I. G., Gajdon, G. K., et al. (2020). Why preen others? predictors of allopreening in parrots and corvids and comparisons to grooming in great apes. Ethology, 126 (2), 207–228.

Noske, S., Beeton, R. J. S., & Jarman, P. (1982). Aspects of the behaviour and ecology of the white cockatoo (’cacatua galerita’) and galah (’c. roseicapilla’) in croplands in north-east new south wales.

Oksanen, J., Blanchet, F. G., Kindt, R., Legendre, P., Minchin, P. R., O’hara, R., Simpson, G. L., Solymos, P., Stevens, M. H. H., Wagner, H., et al. (2013). Package ‘vegan’. Community ecology package, version, 2 (9), 1–295.

OpenStreetMap contributors. (2017). Planet dump retrieved from https://planet.osm.org.

Paradis, E. (2010). Pegas: An r package for population genetics with an integrated–modular approach. Bioinformatics, 26 (3), 419–420.

Parsons, K. M., Durban, J. W., Claridge, D. E., Balcomb, K. C., Noble, L. R., & Thompson, P. M. (2003). Kinship as a basis for alliance formation between male bottlenose dolphins, tursiops truncatus, in the bahamas. Animal Behaviour, 66 (1), 185–194.

Pew, J., Muir, P. H., Wang, J., & Frasier, T. R. (2015). Related: An r package for analysing pairwise relatedness from codominant molecular markers. Molecular ecology resources, 15 (3), 557–561.

Pitcher, T. J., Magurran, A. E., & Allan, J. R. (1986). Size-segregative behaviour in minnow shoals. Journal of Fish Biology, 29, 83–95. https://doi.org/10.1111/j.1095-8649.1986.tb05001.x

Power, D. M. (1966). Agonistic behavior and vocalizations of orange-chinned parakeets in captivity. The Condor, 68 (6), 562–581.

QGIS Development Team. (2009). Qgis geographic information system. Open Source Geospatial Foundation. http://qgis.osgeo.org

R Core Team. (2021). R: A language and environment for statistical computing. R Foundation for Statistical Computing. Vienna, Austria. https://www.R-project.org/

Ruckstuhl, K. E. (2007). Sexual segregation in vertebrates: Proximate and ultimate causes [Publisher: Oxford Academic]. Integrative and Comparative Biology, 47 (2), 245–257. https://doi.org/10.1093/icb/icm030

Schino, G., & Aureli, F. (2010). The relative roles of kinship and reciprocity in explaining primate altruism. Ecology letters, 13 (1), 45–50.

Sharp, S. P., McGowan, A., Wood, M. J., & Hatchwell, B. J. (2005). Learned kin recognition cues in a social bird. Nature, 434 (7037), 1127–1130.

Silk, J. B., Beehner, J. C., Bergman, T. J., Crockford, C., Engh, A. L., Moscovice, L. R., Wittig, R. M., Seyfarth, R. M., & Cheney, D. L. (2009). The benefits of social capital: Close social bonds among female baboons enhance offspring survival [Publisher: Royal Society]. Proceedings of the Royal Society B: Biological Sciences, 276 (1670), 3099–3104. https://doi.org/10.1098/rspb.2009.0681

Silk, J. B., Beehner, J. C., Bergman, T. J., Crockford, C., Engh, A. L., Moscovice, L. R., Wittig, R. M., Seyfarth, R. M., & Cheney, D. L. (2010). Strong and consistent social bonds enhance the longevity of female baboons. Current Biology, 20 (15), 1359–1361. https://doi.org/10.1016/j.cub.2010.05.067

Smith, J. E., Memenis, S. K., & Holekamp, K. E. (2007). Rank-related partner choice in the fission–fusion society of the spotted hyena (crocuta crocuta). Behavioral Ecology and Sociobiology, 61 (5), 753–765. https://doi.org/10.1007/s00265-006-0305-y

Sorato, E., Gullett, P. R., Griffith, S. C., & Russell, A. F. (2012). Effects of predation risk on foraging behaviour and group size: Adaptations in a social cooperative species. Animal Behaviour, 84 (4), 823–834. https://doi.org/10.1016/j.anbehav.2012.07.003

Styche, A. (2000). Distribution and behavioural ecology of the sulphur-crested cockatoo (cacatua galerita l.) in new zealand.

Taylor, S. (2002). On the behavioural ecology and vocal communication of the brown-headed parrot (poicephalus cryptoxanthus) (Doctoral dissertation).

Van Hoof, J. (2000). 16• relationships among non-human primate males: A deductive framework. Primate males: Causes and consequences of variation in group composition. Cambridge University Press, Cambridge, UK, 183–191.

Wahlström, L. K. (1994). The significance of male-male aggression for yearling dispersal in roe deer (capreolus capreolus). Behavioral Ecology and Sociobiology, 35 (6), 409–412.

Waldman, B. (1988). The ecology of kin recognition. Annual review of ecology and systematics, 19 (1), 543–571.

Wang, J. (2011). Coancestry: A program for simulating, estimating and analysing relatedness and inbreeding coefficients. Molecular ecology resources, 11 (1), 141–145.

Whitehead, H. (2008). Analyzing animal societies: Quantitative methods for vertebrate social analysis. University of Chicago Press.

Wilkinson, G. S., Carter, G., Bohn, K. M., Caspers, B., Chaverri, G., Farine, D., Günther, L., Kerth, G., Knörnschild, M., Mayer, F., et al. (2019). Kinship, association, and social complexity in bats. Behavioral Ecology and Sociobiology, 73 (1), 7.

Williams, D. A., & Rabenold, K. N. (2005). Male-biased dispersal, female philopatry, and routes to fitness in a social corvid. Journal of Animal Ecology, 74 (1), 150–159.

Wright, T. F., & Dahlin, C. R. (2018). Vocal dialects in parrots: Patterns and processes of cultural evolution. Emu-Austral Ornithology, 118 (1), 50–66.

Wright, T. F., & Wilkinson, G. S. (2001). Population genetic structure and vocal dialects in an amazon parrot. Proceedings of the Royal Society of London. Series B: Biological Sciences, 268 (1467), 609–616.

Zeus, V. M., Reusch, C., & Kerth, G. (2018). Long-term roosting data reveal a unimodular social network in large fission-fusion society of the colony-living natterer’s bat (myotis nattereri). Behavioral Ecology and Sociobiology, 72 (6), 1–13.

Zheng, X., Levine, D., Shen, J., Gogarten, S. M., Laurie, C., & Weir, B. S. (2012). A high-performance computing toolset for relatedness and principal component analysis of snp data. Bioinformatics, 28 (24), 3326–3328.

